# Role of genetic frustrations in cell reprogramming

**DOI:** 10.1101/2025.01.31.635856

**Authors:** Yuxiang Yao, Jieying Zhu, Wenfei Li, Duanqing Pei

**Affiliations:** Laboratory of Cell Fate Control, School of Life Sciences, Westlake University, Hangzhou 310030, China; Westlake Laboratory of Life Sciences and Biomedicine, Hangzhou 310024, China; CAS Key Laboratory of Regenerative Biology, Center for Cell Lineage and Development, Guangzhou Institutes of Biomedicine and Health, Chinese Academy of Sciences, Guangzhou 510530, China; Department of Physics, National Laboratory of Solid State Microstructure, Nanjing University, Nanjing 210093, China

## Abstract

The cell fate transition is a fundamental characteristic of living organisms. By introducing external perturbations, it is possible to artificially intervene in cell fate and trigger cell reprogramming. Revealing the general principle underlying the induced phenotypic reshaping of cell populations remains a central focus in the field of cell biology. In this study, we investigate the energetic and dynamic features of induced cell phenotypic transition from differentiated somatic state to pluripotent state by constructing a Boolean genetic network model. The simulation and experimental results highlight the critical role of genetic frustration in initiating cell fate transitions, although the two ending phenotypic states are typically featured by minimal frustration. In addition, the altered gene expression profiles exhibit a scale-free distribution, suggesting that there exist a small number of critical genes responsible for the cell fate transition. This study provides important insights into the dynamic principles governing effective cell reprogramming caused by artificial or exogenous interventions.

---

Cell fate transitions occur frequently in organisms, driven by programmed regulation of gene expression at multiple levels [1]. Unveiling the general principles of cell fate transitions is among the key aspects for understanding the nature of life. Typical examples of cell fate transitions include embryonic development and somatic cell reprogramming. Unlike embryonic development, which typically follows well-established roadmaps, somatic cell reprogramming is often subject to extrinsic interference [2]. By simply introducing a few transcription factors (TF) [3–12] under appropriate conditions [13–18], it is possible to convert the cell state of mouse embryonic fibroblasts (MEFs) into that of induced pluripotent stem cells (iPSCs). Breaking existing genetic patterns of cells and rebuilding targeted ones are pivotal tasks for the cell reprogramming.

The canonical Waddington epigenetic landscape paradigm elucidates that genetic circuits, external conditions, and noises collectively determine the valleys and ridges on the genetic topographic maps, which represent cell states and barriers of fate transitions, respectively [19–22]. Current theoretical models of cell reprogramming involving iPSCs can reasonably describe transition trajectories, predict candidate factors, and even propose controlling strategies [23–27]. These theoretical works provide valuable regulatory guidelines for the inducing processes. However, many key aspects on the induced phenotypic reshaping of cell populations is still not fully understood. For instance, why do specific factors contribute to reprogramming rather than others? Is there any common energetic feature in the genetic regulatory network for the productive cell reprogramming events? In a previous work by Font-Clos et al. [28], a Boolean network model based on experimental data was constructed and utilized to explore the cell state transition between epithelial and mesenchymal states. They elucidated high cell phenotypic plasticity with significant population of metastable hybrid cell states along the phenotypic landscape. In a recent work, Tripathi et al. demonstrated that the gene regulation network determining cell fates are minimally frustrated in the steady cell state, which is the key feature distinguishing biological regulatory networks from random networks [29]. More recently, Wang et al. revealed a concerted mechanism of cell phenotypic transitions by analyzing single-cell RNA sequencing (scRNA-seq) data [30]. Inspired by these works, we further explore the energetic and dynamic features essential for successful cell fate transitions by investigating somatic cell reprogramming processes using a Boolean modeling framework of gene regulation. Our numerical results and sequencing data reveal that cells leverage local frustration in their genetic network to facilitate productive fate transitions under external interference, despite the global gene regulation network being minimally frustrated. External interference specifically introduces local frustration to the gene nodes involving activation/inactivation transitions, which triggers the reestablishment of cellular order and ultimately alters cell fates. This local frustration not only decreases the energy barrier height during the fate transition but also shifts the location of the transition state ensemble along the transition pathway [31]. There exists small number of critical genes which are responsible for the cell fate transitions. Our results shed important insights into the general principle of induced cell fate transitions.

## Dynamic framework of cell reprogramming

Experiments reveal that genetic changes during reprogramming typically exhibit a binary-opposite pattern [3–5]. Boolean networks (BNs), which inherently have binary features, can be well served as the dynamic model framework to describe the cell reprogramming. BNs have been utilized in the analysis of biosystems for a long time [32– 34]. In this model, Boolean values 1 and 0 represent the active and silent states of genes, respectively. Consequently, a binary vector 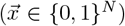 characterizes the state of a genetic network comprising N components. BNs abstract regulatory relations as Boolean functions and network topological structures, effectively catching the essential features of the systems but avoiding parametric explosion. The dynamic rules of system can be described as [35],

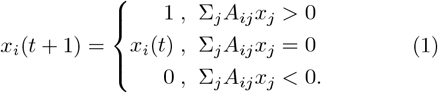

Where A_*ij*_ = ± 1 represents the activation or inhibition operation from gene *j* to gene *i*. The summation runs over all the other genes with a direct regulatory relation with the gene i [Fig. 1(a)]. The system evolution employs an asynchronous rule, wherein only one component is randomly selected to update its state at each time step. 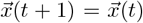 indicates that the system has reached a steady state. The matrix {A_*ij*_}, which describes the regulatory relation of different genes, was extracted from experimental data on somatic cell reprogramming reported in the literature. The gene regulatory network contains 88 genetic components and 387 regulatory edges. More detailed settings and the network structure can be found in the Supplementary Material (SM).

**FIG. 1.**
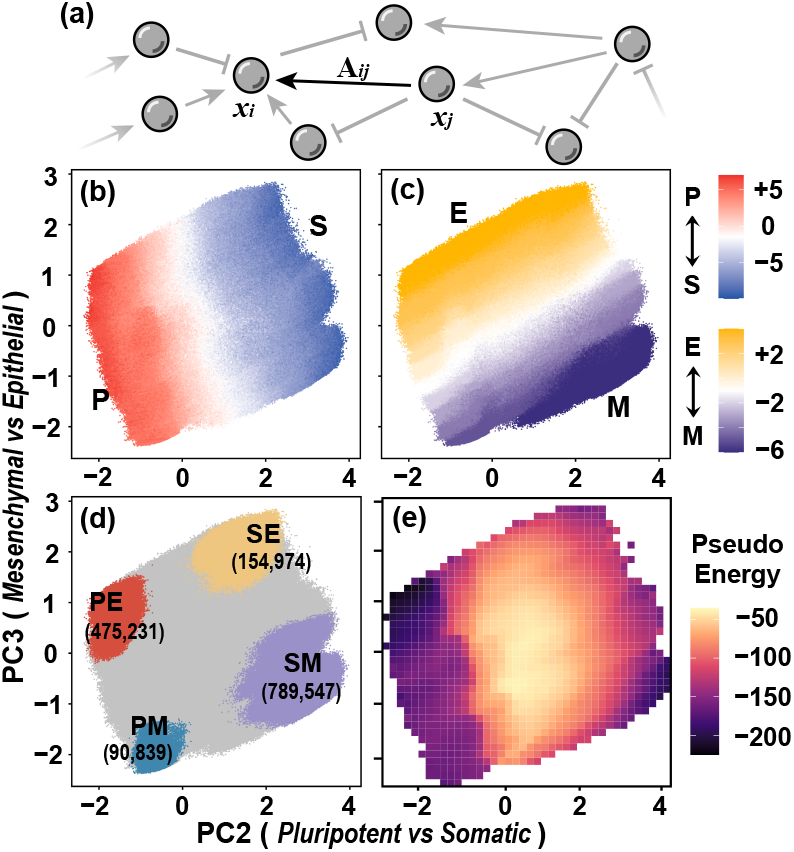
Dynamic framework and phenotypic landscapes of reprogramming systems. (a) Schematic diagram of genetic regulatory. Regulatory patterns include activation (arrow) and inhibition (T-shaped arrow). The arrow A_*ij*_ indicate the regulatory relationship from x_*j*_ to x_*i*_. (b,c) Projection of the obtained 10^7^ steady states along the two essential collective variables PC2 and PC3 from principle component analysis. The steady states are colored according to the phenotypic scores Φ_*PS*_ (b) and Φ_*EM*_ (c). (d) Same as (b,c), but with the steady states colored according to their phenotypic scores, highlighting the top and bottom 20%. (e) Averaged pseudoenergy of the steady states along the two essential collective variables PC2 and PC3.

Cell reprogramming is always accompanied by phenotypic shifts. Induced cells gradually lose their differentiated characteristics and reestablish pluripotent potential, corresponding to a transition from somatic state (S) to pluripotent state (P). Moreover, the cell morphology often undergoes a transition from a mesenchymal-like (M) state to an epithelial-like (E) state as featured by weaker migration capacity. In this study, we characterize the transitions of these two cellular phenotypes together, quantified by two phenotypic scores Φ_*PS*_ and Φ_*EM*_, respectively. The phenotypic scores are defined by

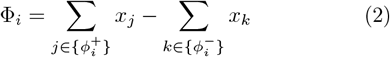

where 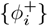 and 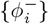 respectively denote positive and negative marker genes of the phenotype *i* (*i*=PS or EM) (See Tab. S1 in SM).

### Genetic energy landscape along two-dimensional essential space

We randomly generate 10^7^ initial states and simulate the relaxation dynamics to the steady states following Eq. 1. On average, the system took 32.7 steps to reach the final steady states (Fig. S1). These steady states collectively form phenotypic phase space that quantify the two phenotypic properties as shown in the principle component analysis [Figs. 1(b,c)]. Cell states located at the peripheral regions along the two-dimensional essential space exhibit the most prominent phenotypic features [Fig. 1(d)]. SM-/PE-like regions respectively act as the source (MEFs) and target (iPSCs) populations of reprogramming processes. In line with previous findings [28, 29], the cell states within each of the four clusters display high similarity, whereas those from different clusters are highly heterogeneous (Fig. S2). These results suggest that successful cell fate transition relies on appropriately breaking and rebuilding cellular order. Therefore, it is crucially important to investigate the underlying dynamic and energetic characteristics of the gene regulatory network in cell fate transitions.

To quantitatively quantify the consistency of the regulatory relationship between different genes, we calculated the cellular pseudo-energy following previous works [28– 30], which is given by

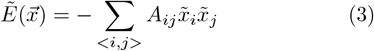

Here 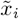 takes the value of +1 and *−*1, respectively, for the active and inactive genes (See Sec. S1.5 in SM). The genetic energy landscape along the two dimensional essential space shows that the cell states with significant phenotypes tend to have lower pseudo-energy [Fig. 1(e)]. This result suggests that these states have well-orchestrated regulatory patterns and therefore low frustration, consistent with previous findings that gene regulatory networks for cell fate decision are minimally frustrated [29]. In addition, the cell states locating between the steady phenotypic states exhibit much higher pseudo energies, which suggests that cell fate transition may involve an activated barrier crossing feature.

## Role of induced genetic frustrations in triggering cell fate transitions

Due to the low frustration of the intrinsic regulatory network, successful cell reprogramming always relies on external perturbations, such as adding specific transcription factors. These external perturbations often lead to the overexpression of specific genes, altering cell fates. Starting from the initial SM states, we performed extensive simulations following the scheme given in Eq. 1. Some trajectories can successfully reach the final PE state, corresponding to productive cell reprogramming events, while others fail to find the final PE state [Figs. 2(a), S3]. By comparing these two sets of trajectories, we may be able to infer the key features contributing to productive cell fate transition. Firstly, we calculated the pseudo-energy distributions for the initial states (sampled from the SM state) of the two sets of trajectories. Interestingly, the initial states for these productive trajectories are featured by much higher pseudo-energies, and therefore higher global frustration, compared to those of the unproductive trajectories. For example, the average pseudo energies are − 95.91 and − 125.47, respectively, for the two sets of initial states [Figs.2(b), S4]. These results suggest that sufficient frustration is essential for the initiation of the productive cell fate transition events.

**FIG. 2.**
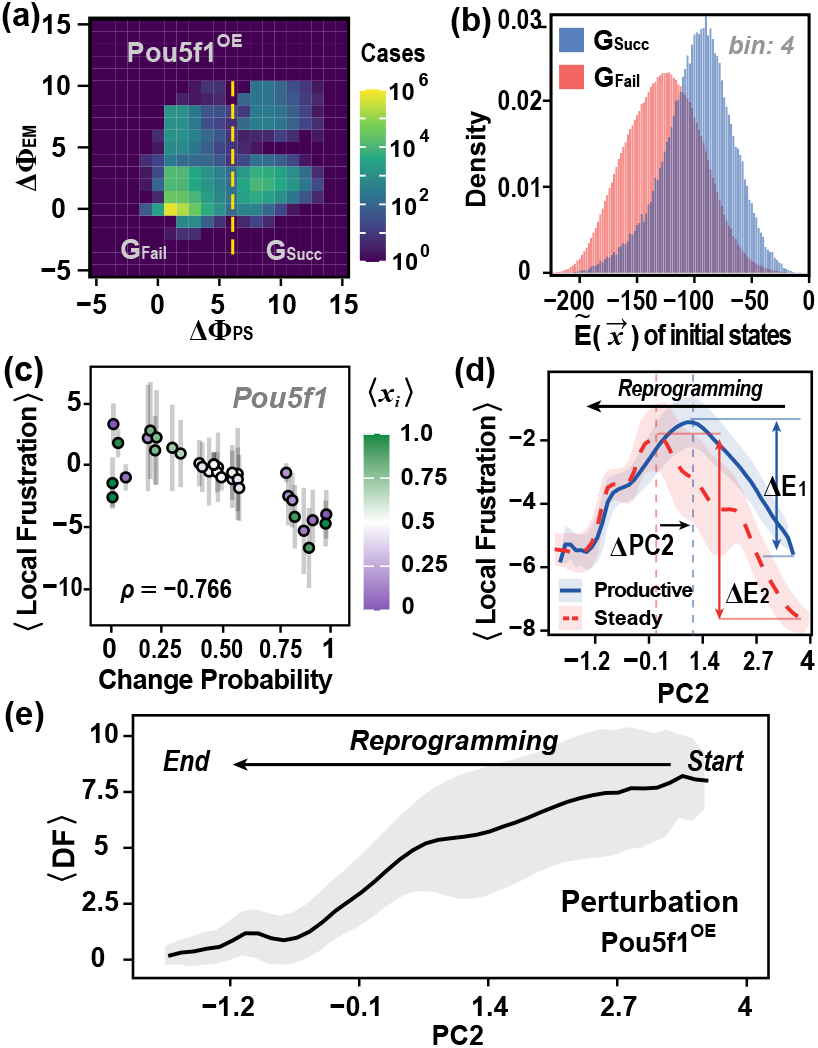
Role of local frustration in perturbation-induced cell fate transition from SM to PE states. (a) Response of MEF-like cells to Pou5f1 overexpression. Simulation trajectories are categorized into two groups: with productive reprogramming (ΔΦ_PS_ > 6) or unproductive reprogramming (ΔΦ_PS_ < 6). The two groups are symbolized as G_Succ_ and G_Fail_, respectively. (b) Distributions of pseudo energies 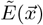 for the initial states of the two groups of simulation trajectories. (c) Correlation between the local frustration of different genes under Pou5f1 overexpression and their probabilities of have different gene states between the initial and final cell states. High-susceptibility genes tend to have lower local frustration. The color of the points represents the average expressions of genes in their initial states. ρ, Pearson correlation. (d) Average local frustration for the high-susceptibility genes along PC2 after introducing Pou5f1 perturbation for the productive reprogramming trajectories (productive). For comparison, the averaged local frustration for the sampled steady states shown in Fig. 1 are also shown (steady). Introducing perturbation elevated the energies of the initial states, decreases the barrier height of cell fate transition, and shifts the location of the transition barrier. ΔE_1_ = 3.11, ΔE_2_ = 4.25, ΔPC2 = 0.65. (e) Average transient driving force ⟨DF⟩ decreases over the course of reprogramming process.

Cell reprogramming is always accompanied with the alteration of activity states of some specific genes. By analyzing the difference of the genetic states between the PE and SM cell states shown in Fig. 1(e), one can identify the susceptible genes subjective to alteration of activity during cell reprogramming (Fig. S5). Genes with a higher probability of changing their states are classified as high-susceptibility genes. In addition to analyzing the feature of global frustration of the gene regulatory network, it is also valuable to investigate the local frustration for each gene in the network. Similar to the pseudoenergies defined above for characterizing global frustration, we can quantify the local frustration by defining the pseudo-energies for individual genes, which is given by 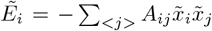. Here, the summation runs over all the N_*i*_ genes regulating gene *i*. Intriguingly, high-susceptibility genes tend to have much lower local pseudo energy compared to other genes in initial cell state, regardless of whether they are expressed. [Figs. 2(c), S6]. Therefore, the low frustration of these susceptible genes can be essential for maintaining the stability of cell states in a noising environment.

We then calculated the single-site pseudo-energy of the high-susceptibility genes for the snapshots sampled in the productive trajectories under Pou5f1 perturbation protocol along the essential reaction coordinate PC2 (Fig. S7). For comparison, we also calculated the single-site pseudoenergies for all the sampled steady states shown in Fig. 1. Similar to the global pseudo-energy landscape shown in Fig. 1(e), the single-site pseudo-energy also exhibits a barrier-crossing feature. The states located around the energy barrier in the energy profile correspond to the rate-limiting transition state ensemble. One can see that pseudo-energies of the snapshots along the reaction coordinate for the productive trajectories not only have higher local frustration at the initial state as discussed above, but also have a lower barrier for the cell reprogramming [Figs. 2(d), S8]. Additionally, the location of the transition state ensemble is shifted towards the initial SM state. These observations also corroborate the findings in Ref. [28], which showed that overexpression of gene SNAI1 leads to a tilt of the phenotypic landscape, favoring the epithelial-mesenchymal transition. These results suggest the crucial role of the induced local genetic frustrations in triggering cell fate transitions. Introducing external perturbations can induce local frustration in the high-susceptibility genes, which in turn reduces the activation barrier of the reprogramming transition and modifies the transition state. Although the overall genetic regulatory network for cell fate decision is minimally frustrated as revealed in previous works [29], local frustration induced by external perturbation is essential for successful cell reprogramming.

The above results are reminiscent of the role of energetic frustration in proteins as elucidated by the energy landscape theory of protein folding [36]. This framework posits that evolution has optimized foldable protein sequences to minimize energetic frustration in their native states, thereby ensuring efficient folding kinetics and structural fidelity. Meanwhile, localized frustration emerges as an evolutionary adaptation at functionally critical regions [37]. Particularly, in many allosteric proteins, for which the functioning dynamics often involve the switching between different conformational states, the residues subjective to conformational motions are often locally frustrated [38–41]. The identified role of localized frustration in governing cell fate transitions within genetic regulatory networks demonstrates remarkable functional congruence with these evolutionarily refined mechanisms in natural protein systems.

The frustration discussed above reflects the inconsistency between gene states and regulatory patterns. With the progression of cell reprogramming, new cellular order needs to re-establish, and the driving force arising from such inconsistency is expected to decrease gradually. To quantitatively quantify such effect, we defined the quantify driving force as following

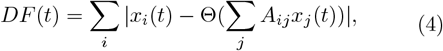

Here, the index *j* runs over all the genes regulating the gene *i*; Θ denotes the dynamic rule given in Eq. 1. As expected, the driving force has a clear decreasing trend as the cell state evolves from the starting somatic state to the final pluripotent state during the reprogramming [Fig. 2(e)].

### Scale-free pattern of the induced cell fate transitions

Yamanaka and other induction protocols utilize a limited number of factors to induce cell reprogramming. Detailed analysis shows that these factors tend to induce more significant local frustration (Fig. S8), as evidenced by substantial changes in single-site pseudo-energy. To assess the order-reshaping potential of each gene, we performed simulations by introducing overexpression (OE) and knockdown (KD) to the genes of the regulatory network. This was realized by setting the genetic states of the corresponding genes to 1 and 0, respectively, throughout the whole simulation trajectories. The simulation starts from a random initial cell state. The results show that OE of different gene has different effect on the variation of the genetic states of the final steady states. To more quantitatively characterize such effect, we calculated the Hamming distance of expression profiles between the initial cell state and the final steady state from the simulations, which is given by 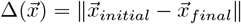, with 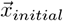 and 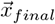 being the genetic states at the initial and final snapshots, respectively, of the simulation trajectory. It corresponds to the total number of genes which have different states at the initial and final cell states. A larger Δ value suggests that the perturbation of a certain gene has a more pronounced effect on the cell fate transition.

Figures 3(a) shows the distribution of Δ from different simulation trajectories. Interestingly, the Δ distribution exhibits a power law pattern over a wide range of Δ values and phenotypic features. Such scale-free patterns suggest that there exist a small number of genes for which OE has the most significant effect on cell phenotypic change. The genes contributing to the heavy tail of the power law distribution may provide potential clues for important regulatory patterns. When examining cases with Δ ⩾ 32, it is observed that these genes predominantly belong to TFs such as Nanog, Sox2, Pou5f1, Esrrb, Klf4, and Sall4, which are associated with maintaining pluripotency [Fig. 3(b)] [2–12]. In addition, the distribution of 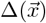 values for these genes tends to have a right-shift, demonstrating the significant effect of these genes on altering cell states (Fig. S9). The results also showed that OE interventions are more effective than KD ones, which is in line with previous experimental observations that overexpressing some TFs instead of suppressing them is more effective in inducing cell reprogramming.

**FIG. 3.**
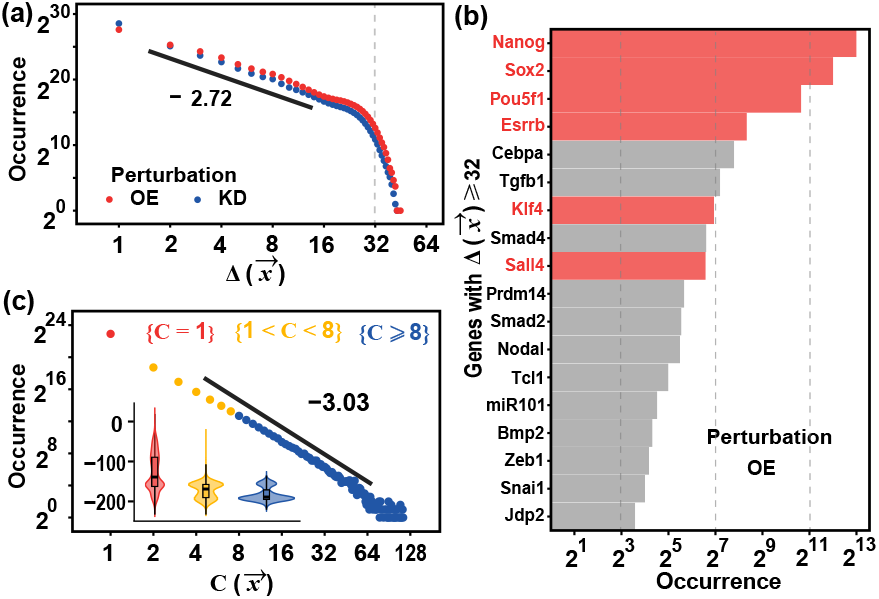
Scale free pattern of the induced cell fate transitions. (a) The distributions of the Hamming distance 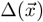 for all the OE/KD simulation trajectories. The simulations are initiated from each of the steady states shown in Fig.1 but with introducing OE (red) or KD (blue) for one of the genes. The repetitive steady states are excluded. (b) Occurrence of the events with 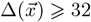 for overexpressing different genes. Only the genes with the occurrence larger than 10 are plotted. Genes marked in red are classic genetic factors commonly used via overexpression in reprogramming experiments. The KD results are presented in Fig. S9. (c) Distribution of the cell-state-capacity 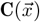 among the steady states shown in Fig. 1. The pseudo energy distributions of the steady states with different 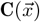 with are also shown as inset.

We next analyzed the features of the final cell states. For each steady state 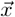, we defined the cell-state-capacity 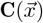, which is given by the number of initial random cell states leading to the final steady state 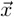 in the simulations. The 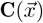 distribution also shows a scaling-free pattern, indicating that majority of the final steady states accommodate only a limited number of initial states. In contrast, a small number of steady state can accommodate a large number of initial states. These large domains of attraction often have lower pseudo energy, indicating that ordered cell states demonstrate stronger dynamic attraction [Fig. 3(c)]. As validation, we numerically simulated the reshaping effects of some key genes on cell phenotypes mentioned in the literature [23, 24, 28]. The results indicate that these genes not only influence the overall phenotype distribution but also, together with the initial cells, determine the extent of phenotype alteration (Fig. S10).

## Experimental validation

As a further demonstration of the crucial role of local frustration in cell fate transition, we analyzed the dynamic and energetic features of mRNA sequencing data (mRNA-Seq) collected in cell reprogramming experiments. In a previous experimental work, Li *et al*. employed the standard Yamanaka protocol to implement reprogramming [3]. As a control, an additional experiment was conducted by introducing overexpression of cJun(cJun^OE^). cJun is a transcription factor that protects the epigenetic state of somatic genes but weakens the epigenetic activity of pluripotent genes. To quantify the time series of the cell fate transition, we binarized the bulk mRNA-Seq and embedded it into the phenotypic diagram. For the canonical Yamanaka protocol, successful cell state transition from MEF-/SM-like state to iPSC-/PE-like state were observed [Fig. 4(a)]. In contrast, the cJun^OE^ cases are limited around the initial points, failing to progress from MEF-/SM-like to iPSC-/PE-like regions. Moreover, continuous phenotypic scores also indicate that intermediate trajectories are blocked by abnormal genetic status caused by cJun^OE^ [Fig. 4(b)].

**FIG. 4.**
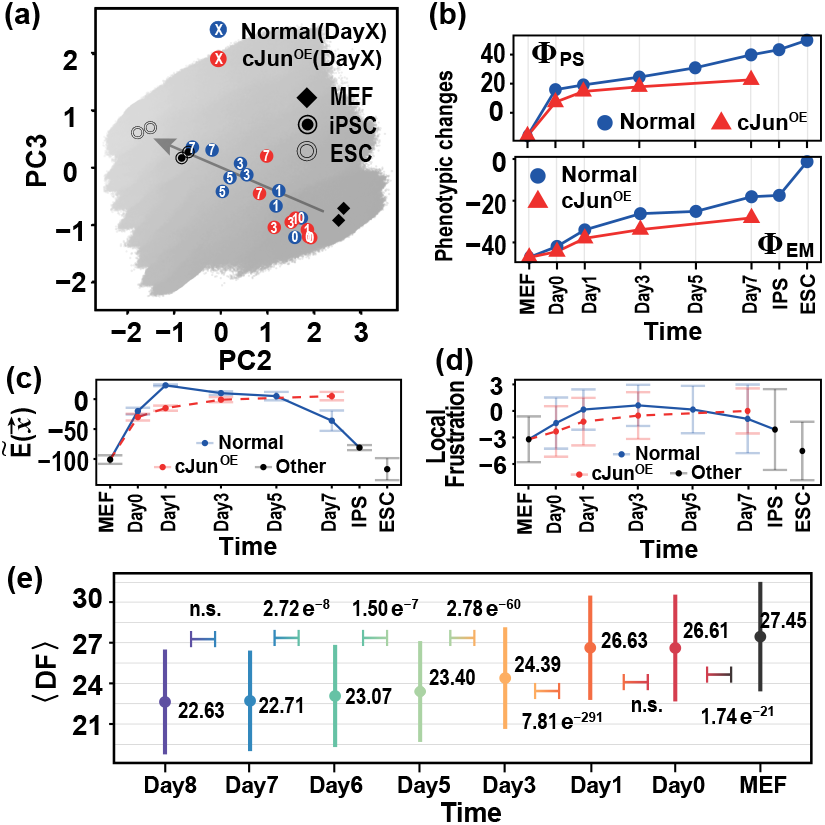
Experimental data showing the changes in various genetic features during somatic cell reprogramming under Yamanaka protocols. (a) Cell reprogramming trajectories from experiments projected along the essential space formed by PC2 and PC3 for the normal Yamanaka protocol and for the Yamanaka protocol with cJun^OE^. MEF, iPSC, and ESC represent mouse embryonic fibroblasts, induced pluripotent stem cells, and embryonic stem cells, respectively. The background landscape is similar as that shown in Fig. 1(b), which only common genes are considered. Note that absolute distances and locations are intended solely as reference, serving to reflect general trends. (b) Phenotypic scores calculated from experimental data along the time series of cell reprogramming with different protocols. The details of the experimental protocols and statistical algorithm can be found in SM Sec 4. (c,d) Pseudo energies (c) and local frustrations (b) along the time series of cell reprogramming for the two experimental protocols. Tag *Other* indicates cells that are not on the reprogramming trajectories, such as MEFs, iPSC, and ESC for reference. (e) Averaged driving force 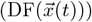 during cell reprogramming from single-cell mRNA-Seq data. The symbols and vertical lines represent the means and standard deviations at each time point. The p-values from t-test showing the significance of the difference between two adjacent distributions are also labeled. To keep consistency with Fig. 2(e), the cell reprogramming progresses from right to left. Details of the distributions are given in Fig. S11.

We then calculated the instantaneous energies of the cell states collected after different time lags in the above two experiments based on the measured genetic activity states. Consistent with the above simulation data, we observed the non-monotonic change of the pseudo energy with the progression of cell reprogramming for the Yamanaka protocol experiment. The pseudo energy increases initially and then decreases over time, which indicates that cell reprogramming involves a energy barrier [Fig. 4(c)]. In contrast, for the cJunOE case, the pseudo energy increases initially and then becomes saturated, lacking an order-reshaping phase essential for cell reprogramming. We also analyzed the local frustrations of various susceptible genes. The local frustration also shows non-monotonic change similar to the time dependence of the global pseudo energy for the Yamanaka protocol [Fig. 4(d)].

To further quantify the role of genetic frustration during the cell reprogramming procedure, we analyzed the distributions of the DF values from the single cell mRNA-Seq with the Yamanaka approach, observing a similar evolutionary pattern as seen in the simulation data of the productive reprogramming events. The mean value of the DF consistently decreases with the progression of the reprogramming [Fig. 4(e); details in Fig. S11], which signifies the reestablishment of the new cellular order towards target cells (Fig. S12).

## Discussion

Cell fate transitions are complex processes orchestrated by established blueprints—regulatory relations. In this study, we consider somatic cell reprogramming processes as paradigmatic cases to capture the key role of genetic frustrations on phenotypic reshaping by performing numerical simulations and cell reprogramming experiments. The results showed that genetic frustrations plays a key role in inducing the cell reprogramming, although the regulatory network are overall minimally frustrated. Critical minority genes within systems contribute to phenotype-oriented perturbations, inducing significant genetic frustrations. In biological scenarios, the extreme inefficiency of transitions indicates inherent obstacles within cells beyond the “blueprints”. The inducing factors, perturbation duration, and epigenetic status collectively complicate the development of universal schemes for controlling cell fate transitions. The fundamental principles of inducing protocols necessitate preparing epigenetic modifications, regularizing transcriptome of starting cells, and creating genetic status of targeted cells [42, 43]. All of these strategies are practically designed to effectively trigger and influence local genetic frustrations. The key role of genetic frustration also underscores the significant genetic inertia of cells, reflecting the role of cellular memory in cell fate decisions [44, 45]. Note that the model frameworks and numerical approaches employed here are preliminary and oversimplified, lacking the complex coupling relations. In future investigations, more precise models, potentially in-corporating other omics [46, 47], are required to provide a clearer understanding of genetic landscapes, thereby more accurately describing cellular states and measuring changes in genetic frustrations. Such an approach will contribute to elucidating the universal principles governing cell fate transitions.

We would like to appreciate Chao Tang for helpful and insightful discussions. Y.Y. and D.P. were supported by Key R&D Program of Zhejiang (2024SSYS0031); W.L. was supported by the National Natural Science Foundation of China (No 11974173); and J.Z. was partially supported by Guangdong Provincial Key Laboratory of Stem Cell and Regenerative Medicine, Guangdong-Hong Kong Joint Laboratory for Stem Cell and Regenerative Medicine.

## References

[1] C. Chen, Y. Gao, W. Liu, and S. Gao, Epigenetic regulation of cell fate transition: learning from early embryo development and somatic cell reprogramming, Biol. Reprod. 107, 183 (2022).

[2] K. Takahashi and S. Yamanaka, Induction of pluripotent stem cells from mouse embryonic and adult fibroblast cultures by defined factors, Cell 126, 663 (2006).

[3] D. Li, J. Liu, X. Yang, C. Zhou, J. Guo, C. Wu, Y. Qin, L. Guo, J. He, S. Yu, H. Liu, X. Wang, F. Wu, J. Kuang, P. Hutchins, J. Chen, and D. Pei, Chromatin accessibility dynamics during iPSC reprogramming, Cell Stem Cell 21, 819 (2017).

[4] B. Wang, L. Wu, D. Li, Y. Liu, J. Guo, C. Li, Y. Yao, Y. Wang, G. Zhao, X. Wang, M. Fu, H. Liu, S. Cao, C. Wu, S. Yu, C. Zhou, Y. Qin, J. Kuang, J. Ming, S. Chu, X. Yang, P. Zhu, G. Pan, J. Chen, J. Liu, and D. Pei, Induction of pluripotent stem cells from mouse embryonic fibroblasts by Jdp2-Jhdm1b-Mkk6-Glis1-Nanog-Essrb-Sall4, Cell Rep. 27, 3473 (2019).

[5] L. Xiao, Z. Huang, Z. Wu, Y. Yang, Z. Zhang, M. Kumar, H. Wu, H. Mao, L. Lin, R. Lin, J. Long, L. Zeng, J. Guo, R. Luo, Y. Li, P. Zhu, B. Liao, L. Wang, and J. Liu, Reconstitution of pluripotency from mouse fibroblast through Sall4 overexpression, Nat. Commun. 15, 10787 (2024).

[6] Y. Li, Q. Zhang, X. Yin, W. Yang, Y. Du, P. Hou, J. Ge, C. Liu, W. Zhang, X. Zhang, Y. Wu, H. Li, K. Liu, C. Wu, Z. Song, Y. Zhao, Y. Shi, and H. Deng, Generation of iPSCs from mouse fibroblasts with a single gene, Oct4, and small molecules, Cell Res. 21, 196 (2011).

[7] L. Guo, L. Lin, X. Wang, M. Gao, S. Cao, Y. Mai, F. Wu, J. Kuang, H. Liu, J. Yang, S. Chu, H. Song, D. Li, Y. Liu, K. Wu, J. Liu, J. Wang, G. Pan, A. P. Hutchins, J. Liu, D. Pei, and J. Chen, Resolving cell fate decisions during somatic cell reprogramming by single-cell RNA-Seq, Mol. Cell 73, 815 (2019).

[8] C. Chronis, P. Fiziev, B. Papp, S. Butz, G. Bonora, S. Sabri, J. Ernst, and K. Plath, Cooperative binding of transcription factors orchestrates reprogramming, Cell 168, 442 (2017).

[9] B. Wang, C. Li, J. Ming, L. Wu, S. Fang, Y. Huang, L. Lin, H. Liu, J. Kuang, C. Zhao, X. Huang, H. Feng, J. Guo, X. Yang, L. Guo, X. Zhang, J. Chen, J. Liu, P. Zhu, and D. Pei, The NuRD complex cooperates with SALL4 to orchestrate reprogramming, Nat. Commun. 14, 2846 (2023).

[10] N. Jain, Y. Goyal, M. C. Dunagin, C. J. Cote, I. A. Mellis, B. Emert, C. L. Jiang, I. P. Dardani, S. Reffsin, M. Arnett, W. Yang, and A. Raj, Retrospective identification of cell-intrinsic factors that mark pluripotency potential in rare somatic cells, Cell Syst. 15, 109 (2024).

[11] V. Malik, L. V. Glaser, D. Zimmer, S. Velychko, M. Weng, M. Holzner, M. Arend, Y. Chen, Y. Srivastava, V. Veerapandian, Z. Shah, M. A. Esteban, H. Wang, J. Chen, H. R. Scholer, A. P. Hutchins, S. H. Meijsing, S. Pott, and R. Jauch, Pluripotency reprogramming by competent and incompetent POU factors uncovers temporal dependency for Oct4 and Sox2, Nat. Commun. 10, 3477 (2019).

[12] H. Benchetrit, M. Jaber, V. Zayat, S. Sebban, Pushett, K. Makedonski, Z. Zakheim, A. Radwan, N. Maoz, R. Lasry, N. Renous, M. Inbar, O. Ram, T. Kaplan, and Y. Buganim, Direct induction of the three preimplantation blastocyst cell types from fibroblasts, Cell Stem Cell 24, 983 (2019).

[13] K. Woltjen and W. L. Stanford, Inhibition of tgf-β signaling improves mouse fibroblast reprogramming, Cell Stem Cell 5, 457 (2009).

[14] B. Feng, J.-H. Ng, J.-C. D. Heng, and H.-H. Ng, Molecules that promote or enhance reprogramming of somatic cells to induced pluripotent stem cells, Cell Stem Cell 4, 301 (2009).

[15] J. Liu, Q. Han, T. Peng, M. Peng, B. Wei, D. Li, X. Wang, S. Yu, J. Yang, S. Cao, K. Huang, A. P. Hutchins, H. Liu, J. Kuang, Z. Zhou, J. Chen, H. Wu, L. Guo, Y. Chen, Y. Chen, X. Li, H. Wu, B. Liao, W. He, H. Song, H. Yao, G. Pan, J. Chen, and D. Pei, The oncogene c-Jun impedes somatic cell reprogramming, Nat. Cell Biol. 17, 856 (2015).

[16] B. D. Stefano, S. Collombet, J. S. Jakobsen, M. Wierer, J. L. Sardina, A. Lackner, R. Stadhouders, C. Segura-Morales, M. Francesconi, F. Limone, M. Mann, B. Porse, D. Thieffry, and T. Graf, C/EBP-α creates elite cells for ipsc reprogramming by upregulating Klf4 and increasing the levels of Lsd1 and Brd4, Nat. Cell Biol. 18, 371 (2016).

[17] P. Hou, Y. Li, X. Zhang, C. Liu, J. Guan, H. Li, T. Zhao, J. Ye, W. Yang, K. Liu, J. Ge, J. Xu, Q. Zhang, Y. Zhao, and H. Deng, Pluripotent stem cells induced from mouse somatic cells by small-molecule compounds, Science 341, 651 (2013).

[18] X. Fu, Q. Zhuang, I. A. Babarinde, L. Shi, G. Ma, H. Hu, Y. Li, J. Chen, Z. Xiao, B. Deng, L. Sun, R. Jauch, and A. P. Hutchins, Restricting epigenetic activity promotes the reprogramming of transformed cells to pluripotency in a line-specific manner, Cell Death Discov. 9, 245 (2023).

[19] C. Waddington, The Strategy of the Genes: a Discussion of Some Aspects of Theoretical Biology (Allen & Unwin, London, 1957).

[20] S. Huang, G. Eichler, Y. Bar-Yam, and D. E. Ingber, Cell fates as high-dimensional attractor states of a complex gene regulatory network, Phys. Rev. Lett. 94, 128701 (2005).

[21] B. Huang, M. Lu, D. Jia, E. Ben-Jacob, H. Levine, and J. N. Onuchic, Interrogating the topological robustness of gene regulatory circuits by randomization, PLoS Comput. Biol. 13, e1005456 (2017).

[22] M. A. Coomer, L. Ham, and M. P. Stumpf, Noise distorts the epigenetic landscape and shapes cell-fate decisions, Cell Syst. 13, 83 (2022).

[23] R. Chang, R. Shoemaker, and W. Wang, Systematic search for recipes to generate induced pluripotent stem cells, PLoS Comput. Biol. 7, e1002300 (2011).

[24] C. Li and J. Wang, Quantifying cell fate decisions for differentiation and reprogramming of a human stem cell network: Landscape and biological paths, PLoS Comput. Biol. 9, e1003165 (2013).

[25] T. Miyamoto, C. Furusawa, and K. Kaneko, Pluripotency, differentiation, and reprogramming: A gene expression dynamics model with epigenetic feedback regulation, PLoS Comput. Biol. 11, e1004476 (2015).

[26] N. Folguera-Blasco, E. Cuyàs, J. A. Menéndez, and T. Alarcón, Epigenetic regulation of cell fate reprogramming in aging and disease: A predictive computational model, PLoS Comput. Biol. 14, e1006052 (2018).

[27] S. Bruno, T. M. Schlaeger, and D. Del Vecchio, Epigenetic OCT4 regulatory network: stochastic analysis of cellular reprogramming, npj Syst. Biol. Appl. 10, 3 (2024).

[28] F. Fontclos, S. Zapperi, and C. A. M. La Porta, Topography of epithelial–mesenchymal plasticity, Proc. Natl. Acad. Sci. USA. 115, 5902 (2018).

[29] S. Tripathi, D. A. Kessler, and H. Levine, Biological networks regulating cell fate choice are minimally frustrated, Phys. Rev. Lett. 125, 088101 (2020).

[30] W. Wang, K. Ni, D. Poe, and J. Xing, Transiently increased coordination in gene regulation during cell phenotypic transitions, PRX Life 2, 043009 (2024).

[31] N. Moris, C. Pina, and A. M. Arias, Transition states and cell fate decisions in epigenetic landscapes, Nat. Rev. Genet. 17, 693 (2016).

[32] S. A. Kauffman, Metabolic stability and epigenesis in randomly constructed genetic nets, J. Theor. Biol. 22, 437 (1969).

[33] S. Kauffman, C. Peterson, B. Samuelsson, and C. Troein, Random Boolean network models and the yeast transcriptional network, Proc. Natl. Acad. Sci. USA. 100, 14796 (2003).

[34] S. A. Kauffman, C. Peterson, B. Samuelsson, and C. Troein, Genetic networks with canalyzing Boolean rules are always stable., Proc. Natl. Acad. Sci. USA. 101, 17102 (2004).

[35] F. Li, T. Long, Y. Lu, Q. Ouyang, and C. Tang, The yeast cell-cycle network is robustly designed, Proc. Natl. Acad. Sci. USA. 101, 4781 (2004).

[36] J. N. Onuchic, Z. Luthey-Schulten, and P. G. Wolynes, Theory of protein folding: the energy landscape perspective, Annu. Rev. Phys. Chem. 48, 545 (1997).

[37] D. U. Ferreiro, E. A. Komives, and P. G. Wolynes, Frustration in biomolecules, Q. Rev. Biophys. 47, 285 (2014).

[38] D. U. Ferreiro, J. A. Hegler, E. A. Komives, and P. G. Wolynes, On the role of frustration in the energy landscapes of allosteric proteins, Proc. Natl. Acad. Sci. USA. 108, 3499 (2011).

[39] W. Li, P. G. Wolynes, and S. Takada, Frustration, specific sequence dependence, and nonlinearity in large-amplitude fluctuations of allosteric proteins, Proc. Natl. Acad. Sci. USA. 108, 3504 (2011).

[40] W. Li, J. Wang, J. Zhang, S. Takada, and W. Wang, Overcoming the bottleneck of the enzymatic cycle by steric frustration, Phys. Rev. Lett. 122, 238102 (2019).

[41] X. Guan, Q.-Y. Tang, W. Ren, M. Chen, W. Wang, P. G. Wolynes, and W. Li, Predicting protein conformational motions using energetic frustration analysis and AlphaFold2, Proc. Natl. Acad. Sci. USA. 121, e2410662121 (2024).

[42] R. Sridharan, M. Gonzales-Cope, C. Chronis, G. Bonora, R. McKee, C. Huang, S. Patel, D. Lopez, N. Mishra, M. Pellegrini, M. Carey, B. A. Garcia, and K. Plath, Proteomic and genomic approaches reveal critical functions of H3K9 methylation and heterochromatin protein1gamma in reprogramming to pluripotency, Nat. Cell Biol. 15, 872 (2013).

[43] A. Mayran and J. Drouin, Pioneer transcription factors shape the epigenetic landscape, J. Biol. Chem. 293, 13795 (2018).

[44] P. S. Stumpf, F. Arai, and B. D. MacArthur, Heterogeneity and ‘memory’ in stem cell populations, Phys. Biol. 17, 065013 (2020).

[45] A. Elsherbiny and G. Dobreva, Epigenetic memory of cell fate commitment, Curr. Opin. Cell Biol. 69, 80 (2021).

[46] L. Li, K. Chen, T. Wang, Y. Wu, G. Xing, M. Chen, Z. Hao, C. Zhang, J. Zhang, B. Ma, Z. Liu, H. Yuan, Z. Liu, Q. Long, Y. Zhou, J. Qi, D. Zhao, M. Gao, D. Pei, J. Nie, D. Ye, G. Pan, and X. Liu, Glis1 facilitates induction of pluripotency via an epigenome-metabolome-epigenome signalling cascade, Nat. Metab. 2, 882 (2020).

[47] W. Li, Q. Long, H. Wu, Y. Zhou, L. Duan, H. Yuan, Y. Ding, Y. Huang, Y. Wu, J. Huang, D. Liu, B. Chen, J. Zhang, J. Qi, S. Du, L. Li, Y. Liu, Z. Ruan, Z. Liu, Z. Liu, Y. Zhao, J. Lu, J. Wang, W.-Y. Chan, and X. Liu, Nuclear localization of mitochondrial TCA cycle enzymes modulates pluripotency via histone acetylation, Nat. Commun. 13, 7414 (2022).

